# Metabolic Modeling of Cystic Fibrosis Airway Communities Predicts Mechanisms of Pathogen Dominance

**DOI:** 10.1101/520619

**Authors:** Michael A. Henson, Giulia Orazi, Poonam Phalak, George A. O’Toole

**Affiliations:** Department of Chemical Engineering and Institute for Applied Life Sciences, University of Massachusetts, Amherst, MA 01003 USA; Department of Microbiology and Immunology, Geisel School of Medicine at Dartmouth, Hanover, NH USA

## Abstract

Cystic fibrosis (CF) is a fatal genetic disease characterized by chronic lung infections due to aberrant mucus production and the inability to clear invading pathogens. The traditional view that CF infections are caused by a single pathogen has been replaced by the realization that the CF lung usually is colonized by a complex community of bacteria, fungi and viruses. To help unravel the complex interplay between the CF lung environment and the infecting microbial community, we developed a community metabolic model comprised of the 17 most abundant bacterial taxa, which account for >95% of reads across samples, from three published studies in which 75 sputum samples from 46 adult CF patients were analyzed by 16S rRNA gene sequencing. The community model was able to correctly predict high abundances of the “rare” pathogens *Enterobacteriaceae, Burkholderia* and *Achromobacter* in three patients whose polymicrobial infections were dominated by these pathogens. With these three pathogens were removed, the model correctly predicted that the remaining 43 patients would be dominated by *Pseudomonas* and/or *Streptococcus*. This dominance was predicted to be driven by relatively high monoculture growth rates of *Pseudomonas* and *Streptococcus* as well as their ability to efficiently consume amino acids, organic acids and alcohols secreted by other community members. Sample-by-sample heterogeneity of community composition could be qualitatively captured through random variation of the simulated metabolic environment, suggesting that experimental studies directly linking CF lung metabolomics and 16S sequencing could provide important insights into disease progression and treatment efficacy.

**Importance:** Cystic fibrosis (CF) is a genetic disease in which chronic airway infections and lung inflammation result in respiratory failure. CF airway infections are usually caused by bacterial communities that are difficult to eradicate with available antibiotics. Using species abundance data for clinically stable adult CF patients assimilated from three published studies, we developed a metabolic model of CF airway communities to better understand the interactions between bacterial species and between the bacterial community and the lung environment. Our model predicted that clinically-observed CF pathogens could establish dominance over other community members across a range of lung nutrient conditions. Heterogeneity of species abundances across 75 patient samples could be predicted by assuming that sample-to-sample heterogeneity was attributable to random variations in the CF nutrient environment. Our model predictions provide new insights into the metabolic determinants of pathogen dominance in the CF lung and could facilitate the development of improved treatment strategies.

## Introduction

Cystic fibrosis is a genetic disease which results in excessive mucus production that reduces lung function and impedes the release of pancreatic enzymes (1, 2). While digestive problems are highly prevalent among CF patients (3), approximately 80-95% of CF deaths are attributable to respiratory failure due to chronic airway infections and associated inflammation (1). The Cystic Fibrosis Foundation (CFF) estimates that approximately 70,000 CF patients are living worldwide and about 1,000 new CF cases are diagnosed in the United States each year (www.cff.org). Following Koch’s postulate (4), the traditional view of CF lung infections has been that specific airway pathogens are responsible for monomicrobial infections (5). CF bacterial pathogens that have been identified from patient sputum samples and commonly studied *in vitro* using pure culture include *Pseudomonas aeruginosa, Haemophilus influenzae, Staphylococcus aureus* and *Burkholderia cepacia* complex, including antibiotic-resistant strains such as methicillin-resistant *S. aureus* (MRSA) and multidrug-resistant *P. aeruginosa* (MDRPA) (1), as well as less common species such as *Achromobacter xylosoxidans, Stenotrophomonas maltophilia* and pathogenic *Escherichia coli* strains (6).

With advent of culture-independent techniques such as 16S rRNA gene amplicon library sequencing, sputum and bronchoscopy samples from CF patients can be analyzed systematically with respect to the diversity and abundance of bacterial taxa present (7, 8). Numerous studies have shown that CF airway infections are rarely monomicrobial, but rather the CF lung harbors a complex community of bacteria that originate from the mouth, skin, intestine and the environment (7-10). 16S sequencing can reliably delineate community members down to the genus level, showing that the most common genera in adult CF patient samples are *Streptococcus, Pseudomonas, Prevotella, Veillonella, Neisseria, Porphorymonas* and *Catonella* (7). While the identities and relative abundances of the genera present can be determined by 16S rRNA gene sequencing, different analysis techniques are required to understand the interactions between the multiple bacterial taxa and the CF lung environment, the role of the individual microbes in shaping community composition and behavior, and the impact of community composition on the efficacy of antibiotic treatment regimens.

*In silico* metabolic modeling has emerged as a powerful approach for analyzing complex microbial communities by integrating genome-scale reconstructions of single-species metabolism within mathematical descriptions of metabolically interacting communities (11, 12). Modeled species interactions typically include competition for host-derived nutrients and cross-feeding of secreted byproducts such as organic acids, alcohols and amino acids between species (13, 14). Due to challenges in developing manually curated reconstructions of poorly studied species, including those present in the CF lung, most *in silico* community models have been restricted to ~5 microbial species (15-17) and fail to adequately cover the diversity of *in vivo* communities. This limitation can be overcome in bacterial communities by using semi-curated reconstructions developed through computational pipelines such as the ModelSeed (18), AGORA (19) and other methods (20). Given the availability of suitable single-strain metabolic reconstructions, a number of alternative methods have been developed for mathematical formulation and numerical solution of microbial community models (21-24). The recently developed SteadyCom method is particularly notable due to its formulation that ensures proper balancing of metabolites across the species and scalability to large communities (25). A properly formulated community model can yield information that is difficult to ascertain experimentally, including the effects of the host environment on community growth, species abundances, and cross-fed metabolite secretion and uptake rates.

In this paper, we utilized 16S rRNA gene amplicon library sequencing data from three published studies (26-28) to develop a 17-species bacterial community model for predicting species abundances in CF airway communities. The 16S rRNA gene sequence data covers 75 distinct sputum samples from 46 adult CF patients, and captures the heterogeneity of CF polymicrobial infections with respect to taxonomic diversity and the prevalence of pathogens including *Pseudomonas, Streptococcus, Burkholderia, Achromobacter* and *Enterobacteriaceae*. The *in silico* community model was used to predict when each pathogen may dominate the polymicrobial infection by using the 16S rRNA gene sequence data to restrict which pathogens were present in the simulated community. By randomly varying the availability of host-derived nutrients, the model was used to simulate sample-by-sample heterogeneity of community compositions across patients and to understand how metabolite cross-feeding enhanced pathogen abundances. To our knowledge, this study represents the first attempt to metabolically model the CF airway bacterial community rather than model the individual metabolism of common CF pathogens (29-34). Furthermore, our approach of directly predicting species abundances rather than using measured abundances as model input data to constrain predictions distinguished our study from other community modeling efforts driven by 16S rRNA gene sequence data (14, 35-37).

## Results

### Few Taxonomic Groups Dominate the CF Airway Community Samples

Principal component analysis (PCA) was performed on the normalized read data of the 75 samples to evaluate sample heterogeneity. The first three principal components (PCs) captured 77.8% of the data variance, with the first PC capturing 57.3% of variance and most heavily weighting the most abundant genera *Pseudomonas, Streptococcus and Prevotella* as expected (Table S5). A considerable degree of heterogeneity was evident from a plot of the 75 samples in the coordinates defined by the first three PCs (Figure 2A). Most striking were the outlier samples from three patients infected with *Enterobacteriaceae* (samples 25-27), *Burkholderia* (samples 19-21) or *Achromobacter* (samples 31, 32) compared to the patients lacking these three organisms (*i.e*., the remaining 67 samples).

**Figure 1.**
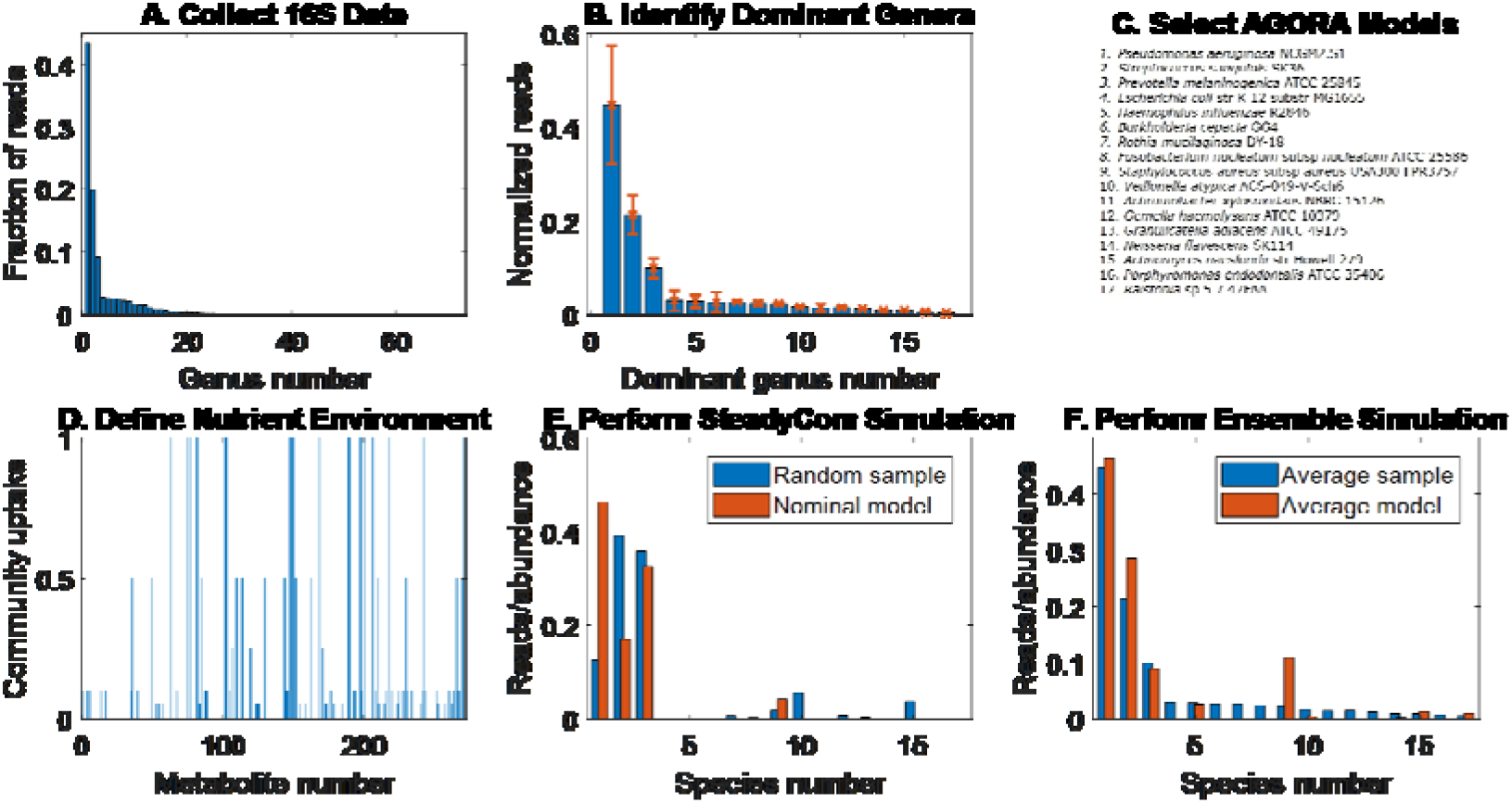
Overview of the community metabolic modeling framework driven by patient microbiota composition data. (A) 16S rRNA gene sequence data for 46 patients averaged across 75 distinct samples for the 72 highest ranked taxonomic groups (typically genera). (B) 16S rRNA gene sequence data for the 17 highest ranked taxonomic groups normalized to sum to unity and then averaged across the 75 samples. The error bars represent the variances of the normalized read data. (C) AGORA strain models (19) selected for 17 species that represent each taxonomic group. (D) Definition of the nutrient environment through specification of the community uptake rate of each extracellular metabolite. (E) Species abundances predicted from a SteadyCom (25) simulation with nominal community uptake rates compared to normalized reads for a random patient sample. (F) Average species abundances predicted from an ensemble of SteadyCom simulations with randomized community uptake rates compared to normalized reads averaged across the patient samples.

**Figure 2.**
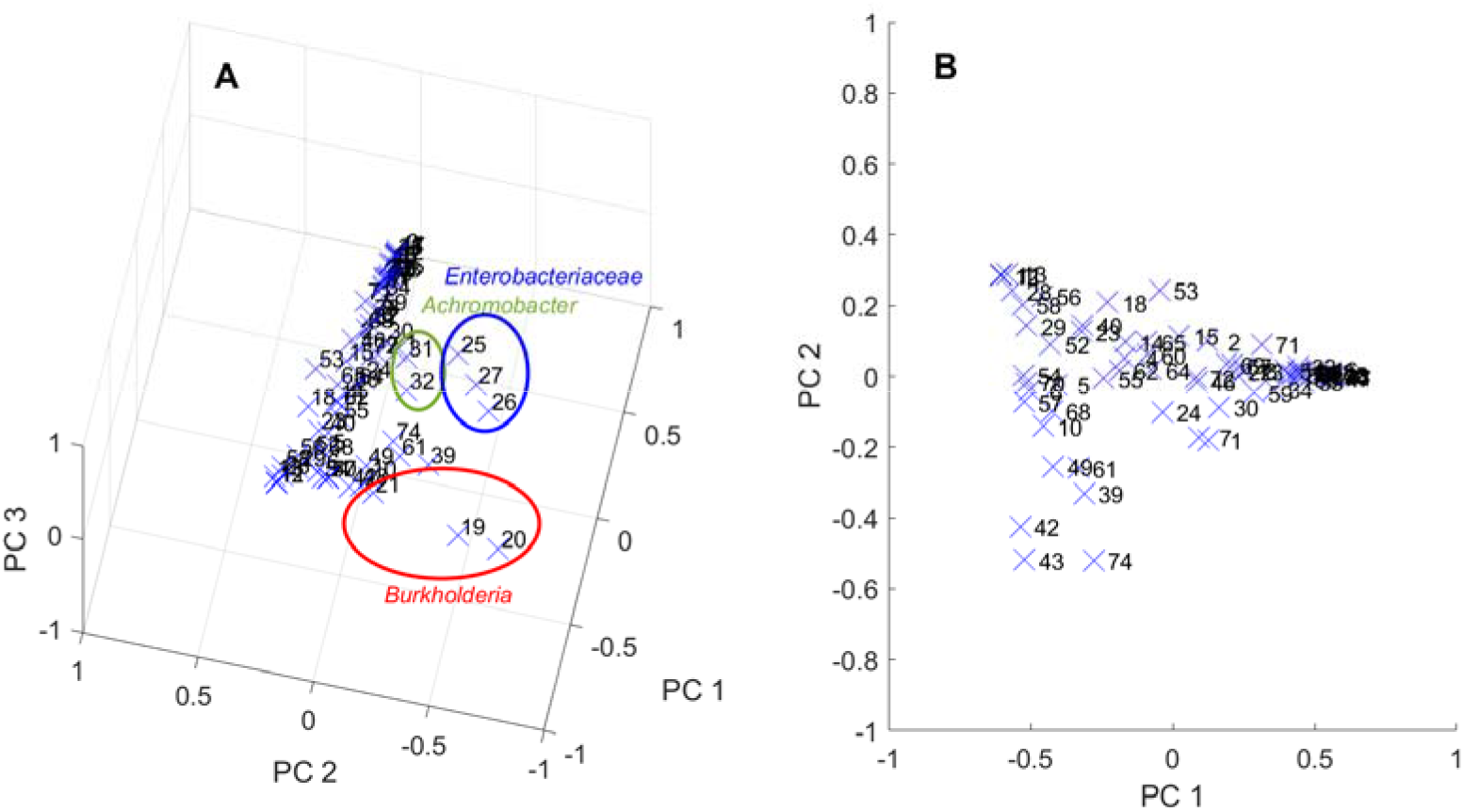
PCA performed on the normalized read data. (A) PCA performed for all 75 samples with the normalized reads for each taxonomic group plotted using the first three principle components (PCs) that explained 57.3%, 12.3% and 8.2%, respectively, of the data variance. Sample points for *Enterobacteriaceae, Burkholderia* and *Achromobacter* appeared as outliers. (B) PCA performed for 67 samples when the 8 samples containing *Enterobacteriaceae, Burkholderia* and *Achromobacter* were removed. The normalized reads for each taxonomic group were plotted using the first two PCs that explained 72.6%, and 11.7%, respectively, of the data variance.

Because each pathogen infected only a single patient among the 46 included patients, we generated a smaller dataset of 67 samples by removing these 8 samples. When PCA was performed on this reduced dataset, the first three PCs explained 92.6% of the data variance (Table S6), suggesting substantially reduced heterogeneity compared to the full dataset. These three PCs heavily weighted only the four taxonomic groups *Pseudomonas, Streptococcus, Prevotella* and *Haemophilus*, with the first PC representing high *Pseudomonas* and low *Streptococcus*, the second PC component representing high *Streptococcus* and moderate *Pseudomonas*, and the third PC representing high *Haemophilus*, low *Pseudomonas* and low *Streptococcus*. Considerable heterogeneity was evident when the 67 samples were plotted using the first two PCs accounting for 84.2% of the variance (Figure 2B). Here the first PC represented high *Pseudomonas*, low *Streptococcus*, moderate *Prevotella* and moderate *Haemophilus*, and the second PC represented low *Pseudomonas*, high *Streptococcus*, low *Prevotella* and low *Haemophilus*.

Based on these results, we focused our community modeling efforts on predicting the infrequent dominance of the pathogens *Enterobacteriaceae*, *Burkholderia* and *Achromobacter*, and the heterogeneity in the abundances of *Pseudomonas, Streptococcus, Prevotella* and *Haemophilus* across the remaining samples. *Pseudomonas*, *Streptococcus* and *Prevotella* have been found by directly sampling the lung of CF patients via bronchoalveolar lavage (38), while *Haemophilus* is a widely-accepted CF pathogen (7). The other 10 genera (Table 1) were maintained in the model to simulate competition/cooperation with the more dominant species and to determine if the relatively low abundances of these genera could be predicted.

**Table 1.** CF genera analyzed. Shown is a list of the 17 species/strains included in the CF airway community model, the normalized fractional reads for the associated genera averaged across the 75 samples, and the percentage of samples in which the normalized reads exceeded 1%.

| Species Number | Species Strain Name | Average Reads | Sample Reads > 1% |
| --- | --- | --- | --- |
| 1 | *Pseudomonas aeruginosa* NCGM2.S1 | 0.447 | 85.3% |
| 2 | *Streptococcus sanguinis* SK36 | 0.213 | 88.0% |
| 3 | *Prevotella melaninogenica* ATCC 25845 | 0.098 | 74.7% |
| 4 | *Escherichia coli* str. K-12 substr. MG1655 | 0.029 | 4.0% |
| 5 | *Haemophilus influenzae* R2846 | 0.028 | 22.7% |
| 6 | *Burkholderia cepacia* GG4 | 0.026 | 4.0% |
| 7 | *Rothia mucilaginosa* DY-18 | 0.026 | 48.0% |
| 8 | *Fusobacterium nucleatum* subsp. *nucleatum* ATCC 25586 | 0.023 | 26.7% |
| 9 | *Staphylococcus aureus* subsp. *aureus* USA300 FPR3757 | 0.023 | 34.7% |
| 10 | *Veillonella atypica* ACS-049-V-Sch6 | 0.016 | 48.0% |
| 11 | *Achromobacter xylosoxidans* NBRC 15126 | 0.014 | 2.7% |
| 12 | *Gemella haemolysans* ATCC 10379 | 0.015 | 30.7% |
| 13 | *Granulicatella adiacens* ATCC 49175 | 0.012 | 36.0% |
| 14 | *Neisseria flavescens* SK114 | 0.008 | 18.7% |
| 15 | *Actinomyces naeslundii* str. Howell 279 | 0.009 | 21.3% |
| 16 | *Porphyromonas endodontalis* ATCC 35406 | 0.006 | 20.0% |
| 17 | *Ralstonia* sp 5 7 47FAA | 0.004 | 6.7% |

### The Community Model Can Reproduce Dominance of CF Pathogens

We simulated the growth of each species individually to compare their monoculture growth rates with the nominal community nutrient uptake rates (Table S4). Interestingly, the three highest growth rates belonged to the rare pathogens *Escherichia*, *Burkholderia* and *Achromobacter*, while the next three highest growth rates belonged to the common pathogens *Pseudomonas*, *Streptococcus* and *Staphylococcus* (Figure 3A; species numbered as in Table 1). These predictions were consistent with our modeling results for the gut microbiome (39) where opportunistic pathogens consistently had higher growth rates than commensal species. The other two species *Prevotella* and *Haemophilus* commonly observed in the 75 patient samples were predicted to have much lower *in silico* growth rates. The three species representing *Fusobacterium, Granulicatella* and *Porphyromonas* did not grow individually due to their inability to meet the defined ATP maintenance demand, although they could grow when strategically combined with other modeled species. For example, *Fusobacterium, Granulicatella* and *Porphyromonas* were predicted to grow in coculture with *Ralstonia, Prevotella* and *Actinomyces*, respectively. The species abundances predicted for a specified nutrient condition depended both on the monoculture growth rates and the ability of each species to efficiently utilized secreted metabolites to enhance its growth rate. These emergent cross-feeding relationships allowed otherwise slower growing species to coexist with species that exhibited high monoculture growth rates.

**Figure 3.**
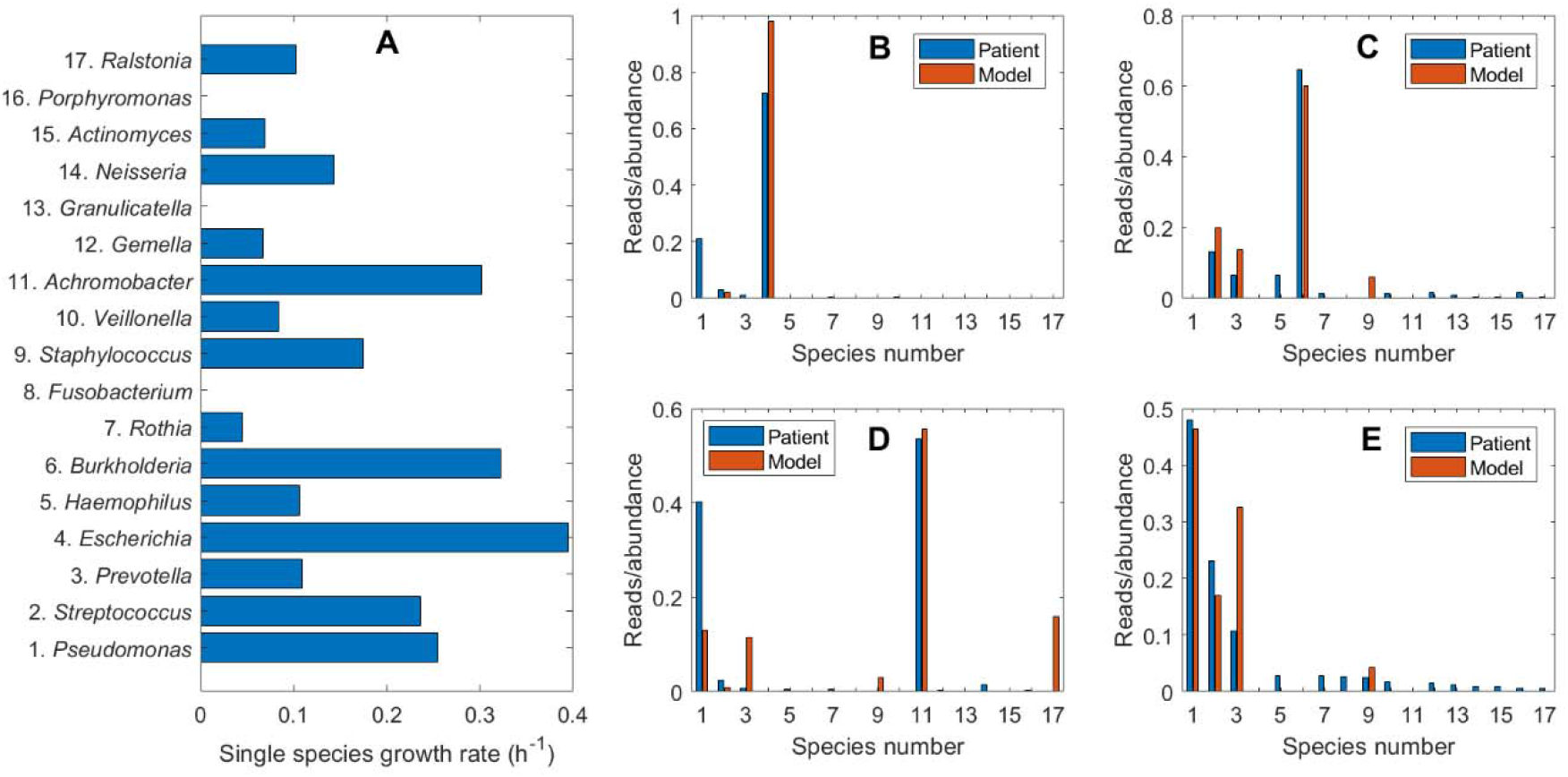
Single-species and community simulations performed with the nominal nutrient uptake rates in Table S3. (A) Single-species growth rates with the species numbered according to Table 1. (B) Comparison of predicted species abundances to the average of the normalized reads for the single patient infected with *Enterobacteriaceae/Escherichia* (samples 25-27). (C) Comparison of predicted species abundances to the average of the normalized reads for the single patient infected with *Burkholderia* (samples 19-21). (D) Comparison of predicted species abundances to the average of the normalized reads for the single patient infected with *Achromobacter* (samples 31, 32). (E) Comparison of predicted species abundances to the average of the normalized reads for the 43 patients not infected with *Enterobacteriaceae/Escherichia, Burkholderia* or *Achromobacter* (samples 1-18, 22-24, 28-30, 33-75).

We conducted simulations using the nominal nutrient uptake rates (Table S4) to determine if the community model could capture dominance of each rare pathogen in the absence of the other two rare pathogens. Each simulation was performed by constraining the abundances of the other two pathogens to zero, effectively producing reduced communities of 15 species. The predicted abundances from each simulation were compared to the normalized reads averaged over the patient samples which contained the associated pathogen: *Enterobacteriaceae/Escherichia* (samples 25-27; Figure 3B), *Burkholderia* (samples 19-21; Figure 3C) or *Achromobacter* (samples 31 and 32; Figure 3D). For each simulated case, the model correctly predicted dominance of the associated pathogen. For the *Burkholderia-* and *Achromobacter-* infected patients, the abundances of the dominant pathogen as well as less prevalent species were well predicted.

We performed simulations for the remaining 43 patients by reducing the community to 14 species by constraining the abundances of all three rare pathogens to zero. The model predicted abundances were compared to the normalized reads averaged over the 67 samples remaining when the 8 rare pathogen-containing samples were removed (Figure 3E). The model correctly predicted that *Pseudomonas, Streptococcus* and *Prevotella* would dominate the community, although the *Prevotella* abundance was overpredicted at the expense of *Streptococcus* as well as several less abundant genera. The only other genus present in the simulated community was *Staphylococcus*, while the averaged reads showed a greater amount of diversity. Compared to the averaged data, individual samples showed less diversity, which is more consistent with model predictions as discussed below.

### The Community Model Can Reproduce Pathogen Heterogeneity Across Airway Samples

The CF airway communities exhibited a substantial degree of sample-to-sample heterogeneity when rare pathogens were present (Figure 2A) or absent (Figure 2B). We performed simulations to assess the extent to which sample-to-sample differences in taxonomic group reads could be explained by heterogeneity in the metabolic environment of the CF lung. More specifically, we randomized the community nutrient uptake rates around their nominal values (Materials and Methods; Table S4) to mimic heterogeneous lung environments shown to occur across CF patients (40, 41) and in longitudinal samples from a single patient (42). Each simulation with a set of randomized uptake rates was termed a “simulated sample,” and we tested the hypothesis that the experimental samples could be interpreted as having been drawn from the much larger set of simulated samples we generated. Due to the relatively small number of *Enterobacteriaceae/Escherichia-, Burkholderia-* and *Achromobacter-containing* samples, we only performed 100 randomized community simulations for each of these pathogens. By contrast, 1000 randomized simulations were performed for communities without these three rare pathogens since the associated patient sample size was comparatively large. The single model simulation that best represented a particular patient sample was determined by the minimum least-squares error between the normalized measured reads and the predicted abundances across all simulations. For the 8 rare pathogen-containing samples, we plotted the measured reads and predicted abundances of the five most common genera (*Pseudomonas, Streptococcus, Prevotella, Haemophilus, Staphylococcus*) and the pathogen of interest. For the remaining 67 samples, we plotted the five most common genera plus the next most abundant genus according to measured reads.

Randomized nutrient simulations were able to generate model predictions that reproduced the major features of the 3 *Enterobacteriaceae/Escherichia-containing* samples (Figure 4A), including the high *Enterobacteriaceae/Escherichia* reads and presence of the other main community members (*Pseudomonas, Streptococcus* and *Prevotella*). The *Streptococcus* reads were predicted relatively accurately, while *Pseudomonas* reads were underpredicted and *Prevotella* reads were overpredicted. As measured by the least-squares error, improved predictions were obtained for the 3 *Burkholderia*-containing samples (Figure 4B). The *Burkholderia* reads were accurately reproduced and *Streptococcus* was correctly predicted to be the second-most abundant genus, suggesting a synergism between these two genera. This prediction has experimental support from *in vitro* experiments showing that mucin-degrading anaerobes such as *Streptococci* promote the growth of CF pathogens such as *B. cenocepacia* when mucins are provided as the sole carbon source (43). The two *Achromobacter-containing* samples were well predicted in terms of *Achromobacter* reads and *Pseudomonas* being the other dominant genus (Figure 4C). These predictions are consistent with an *in vitro* study showing that *Achromobacter* sp. enhanced the ability of multiple *P. aeruginosa* strains to form biofilms (44). Furthermore, a clinical study with 53 patients having positive cultures for *A. xylosoxidans* showed that all 6 patients that were chronically infected by *A. xylosoxidans* were co-infected with *P. aeruginosa* (45). Complete comparisons of the normalized measured reads and model predicted abundances for the 8 samples with the rare pathogens are presented in Table S7, which shows that the model generally produced less diverse communities as measured by the richness (number of species with abundances exceeding 1%) and the equitability (the inverse Simpson metric; (46)).

**Figure 4.**
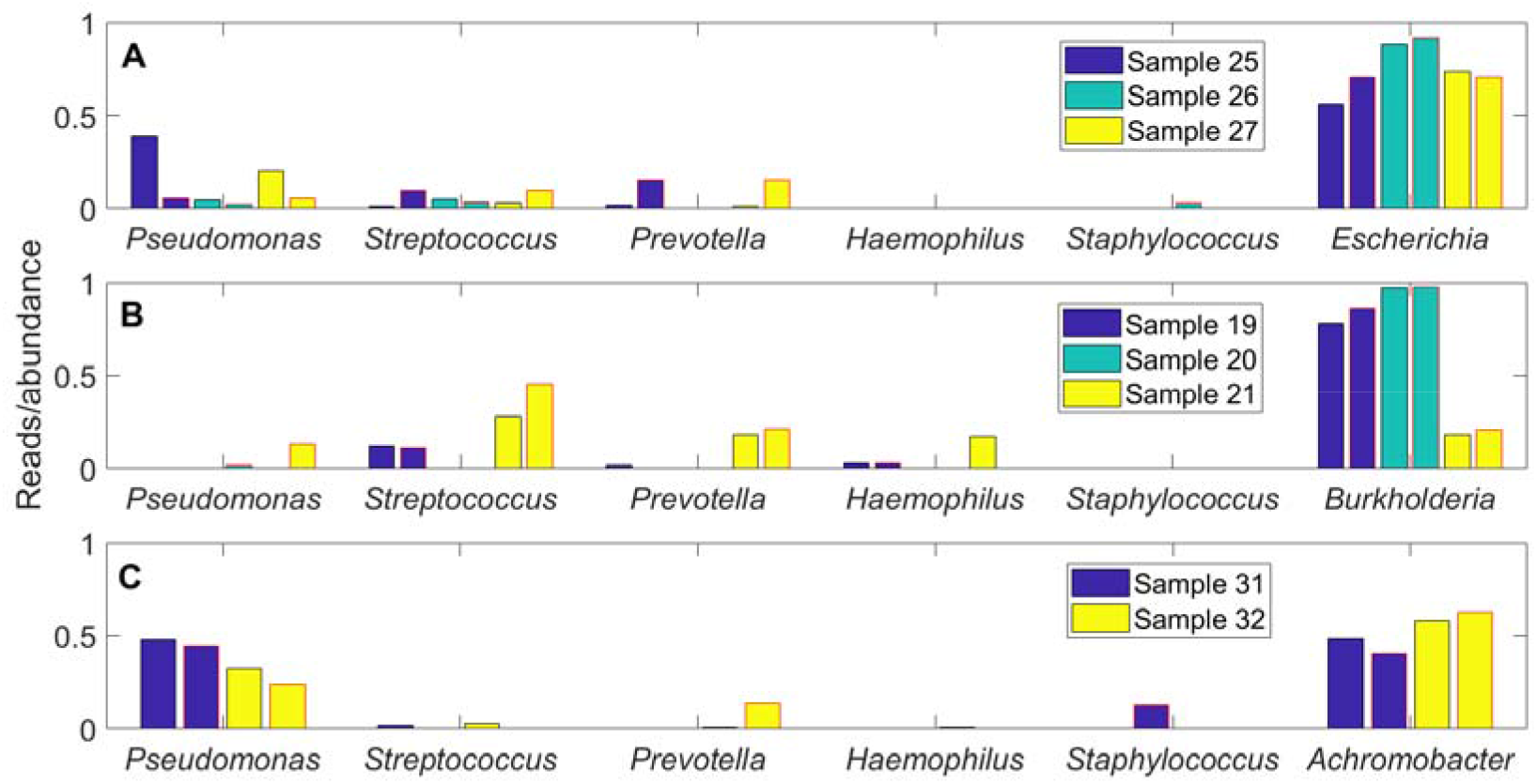
Taxonomic reads for patient samples containing rare pathogens compared to species abundances predicted from community models with randomized nutrient uptake rates. The genera *Pseudomonas, Streptococcus, Prevotella, Haemophilus* and *Staphylococcus* and the indicated rare pathogen (*Enterobacteriaceae/Escherichia, Burkholderia* or *Achromobacter*) are shown for each case. (A) Individual models that best fit the 3 *Enterobacteriaceae/Escherichia*-containing samples 25-27 selected from an ensemble of 100 15-species models without *Burkholderia* or *Achromobacter*. (B) Individual models that best fit the 3 *Burkholderia*-containing samples 19-21 selected from an ensemble of 100 15-species models without *Enterobacteriaceae/Escherichia* or *Achromobacter*. (C) Individual models that best fit the 2 *Achromobacter-containing* samples 31 and 32 selected from an ensemble of 100 15-species models without *Enterobacteriaceae/Escherichia* or *Burkholderia*. Each abundance for a patient sample is shown in the first bar and each abundance predicted by the corresponding model is shown in the second bar with red outline.

The lack of patient samples containing *Enterobacteriaceae/Escherichia, Burkholderia* and *Achromobacter* limited our ability to analyze heterogeneity of communities with these pathogens. By contrast, the 67 samples remaining when the 8 samples containing these three pathogens were removed offered a much larger dataset for heterogeneity analysis. Each of these 67 samples was matched to one of the 1000 randomized model simulations according to the smallest least-squares error between the normalized reads of the sample and the predicted abundances of the model (Table S8). Representative results are shown for patient samples with relatively small (Figure 5A), moderate (Figure 5B) and large (Figure 5C) error values. Samples which were most accurately reproduced generally contained high *Pseudomonas* reads (84%+/-15%) with the remainder of the community consisting of *Streptococcus* and *Prevotella* (Figure 5A). These 22 samples were best matched by 11 distinct models, suggesting that patient samples dominated by *Pseudomonas* contained a higher degree of heterogeneity than the simulated samples.

**Figure 5.**
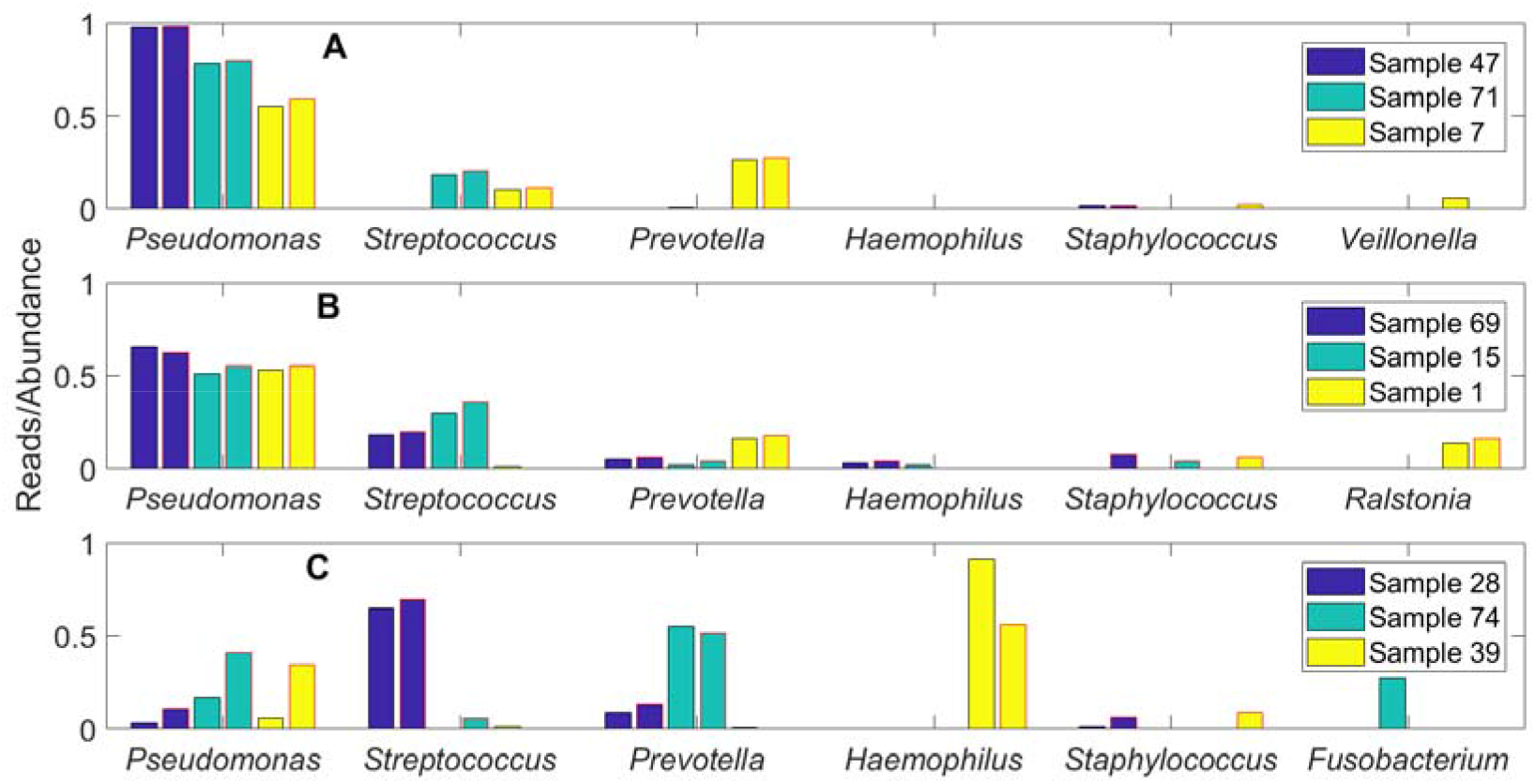
Taxonomic reads for patient samples without rare pathogens compared to species abundances predicted from community models with randomized nutrient uptake rates. The genera *Pseudomonas, Streptococcus, Prevotella, Haemophilus* and *Staphylococcus* and the next most abundant genera are shown for each case. Individual models that best fit the 67 patient samples were selected from an ensemble of 1000 14-species models without *Enterobacteriaceae/Escherichia, Burkholderia* or *Achromobacter*. (A) Three representative samples for which the least-squares error measures were within the smallest third of all samples. (B) Three representative samples for which the least-squares error measures were within the middle third of all samples. (C) Three representative samples for which the least-squares error measures were within the largest third of all samples. Each abundance for a patient sample is shown in the first bar and each abundance predicted by the corresponding model is shown in the second bar with red outline.

The 22 samples which produced moderate prediction errors were characterized by lower and more variable *Pseudomonas* reads (48%+/-28%) as well as more variable distributions of *Streptococcus* and *Prevotella* reads (Figure 5B). The ensemble of randomized models could capture the relative amounts of these three genera, but often predicted the presence of *Staphylococcus* not observed in the patient samples. This discrepancy could be attributable to the unmodeled ability of *Pseudomonas* to secrete diffusible toxins which inhibit *Staphylococcus* respiration and render *Staphylococcus* less metabolically competitive in partially aerobic environments (47) such as the CF lung. Interestingly, the model ensemble could reproduce the relatively high *Ralstonia* reads in sample 1 while also predicting no *Ralstonia* in samples 15 and 69. The 23 samples which produced the largest prediction errors were characterized by much lower *Pseudomonas* reads (13%), higher reads of *Streptococcus* and *Prevotella* (34% and 19%, respectively; e.g. samples 26 and 74 in Figure 5C) and higher representation of less common genera. These samples also produced higher *Haemophilus* reads, primarily due to two *Haemophilus-dominated* samples (e.g. sample 39 in Figure 5C). While the model ensemble generally was able to reproduce the observed *Streptococcus* and *Prevotella* reads in these samples, the models tended to overpredict *Pseudomonas* and *Staphylococcus* at the expense of the less common genera. In particular, the ensemble underpredicted the abundances of *Rothia*, *Fusobacterium* and *Gemella* while the average reads of these three genera across the 23 samples summed to 16% This discrepancy could suggest that these 23 samples were obtained from patients with less advanced CF lung disease, which correlates to higher diversity communities *in vivo* (28, 48).

To gain further insights into the ability of the community model to mimic sample-to-sample heterogeneity in the absence of rare pathogens, we compared read data and abundance predictions in the PC space calculated from the 67 patient samples. Each of the 1000 model simulations was mapped into the two-dimensional space defined by the first two PCs (Figure 2B), which explained 84.2% of normalized read data variance (Table S6). The model ensemble was able to reproduce most of the observed variability as reflected by the cloud of model simulations overlapping most of the patient samples (Figure 6A). The patient and simulated samples covered the same range of the first PC, which was heavily weighted by *Pseudomonas, Streptococcus* and *Prevotella* (Table S6). Importantly, this consistency shows that heterogeneity across these three dominant genera could be predicted from variations in the CF lung metabolic environment, as we hypothesized.

**Figure 6.**
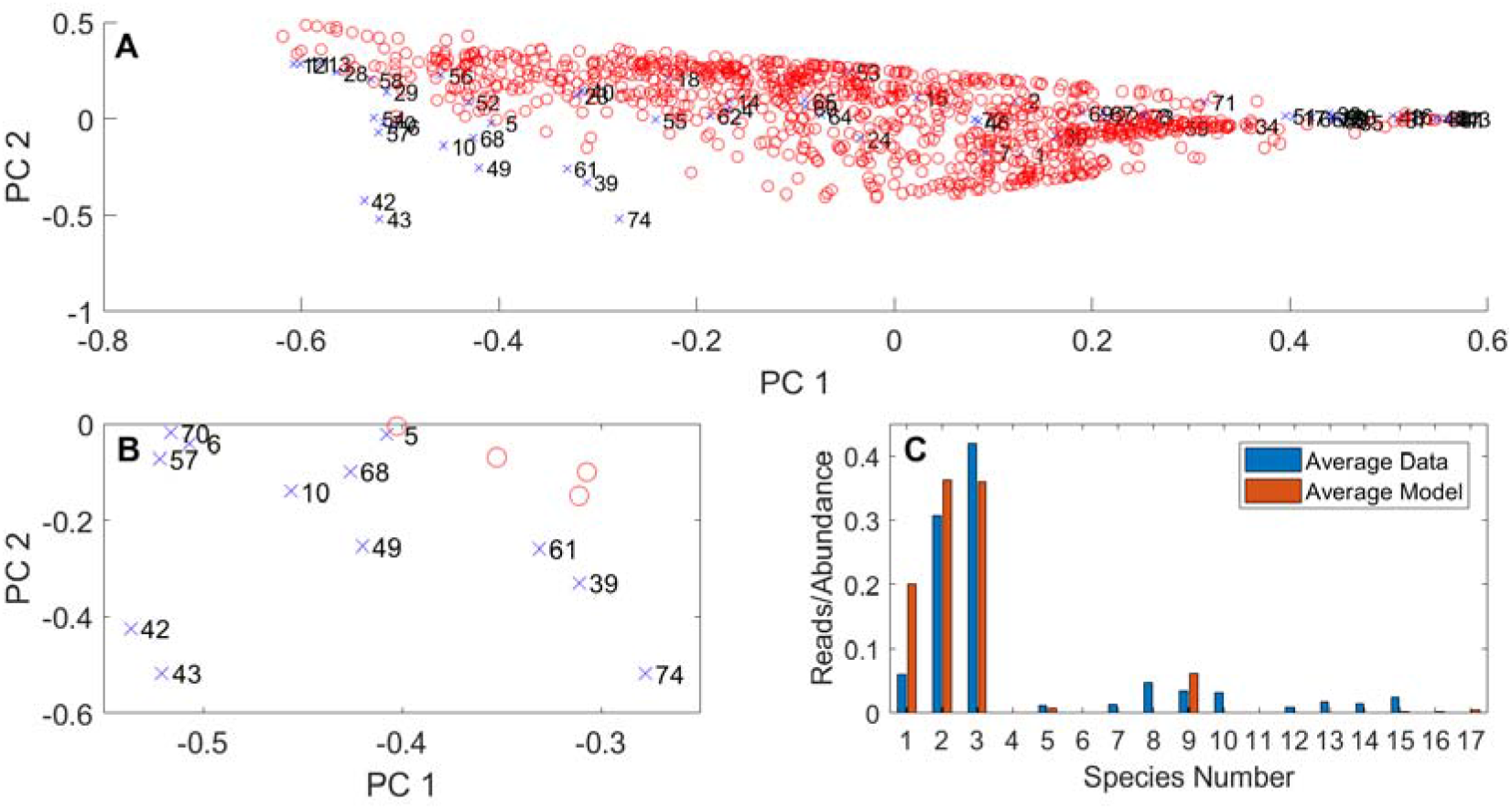
Principal component analysis (PCA) of taxonomic reads for patient samples without rare pathogens and species abundances predicted from 14-species community models with randomized nutrient uptake rates. (A) Representation of the 67 patient samples in the two-dimensional space defined by the first two principal components (PCs) obtained when PCA is performed on the normalized reads of these patient samples. Species abundances predicted from an ensemble of 1000 models transformed into the PC space of the normalized read data. (B) Enlarged view of the lower left portion of the PCA plot in Figure 6A. (C) Average genera reads obtained for 8 samples (5, 6, 10, 39, 42, 43, 49, 57, 61, 68, 70, 74) in Figure 6B with elevated *Prevotella* representation compared to the average abundances predicted from the best-fit models for these 8 samples with the species number as in Table 1.

**Figure 7.**
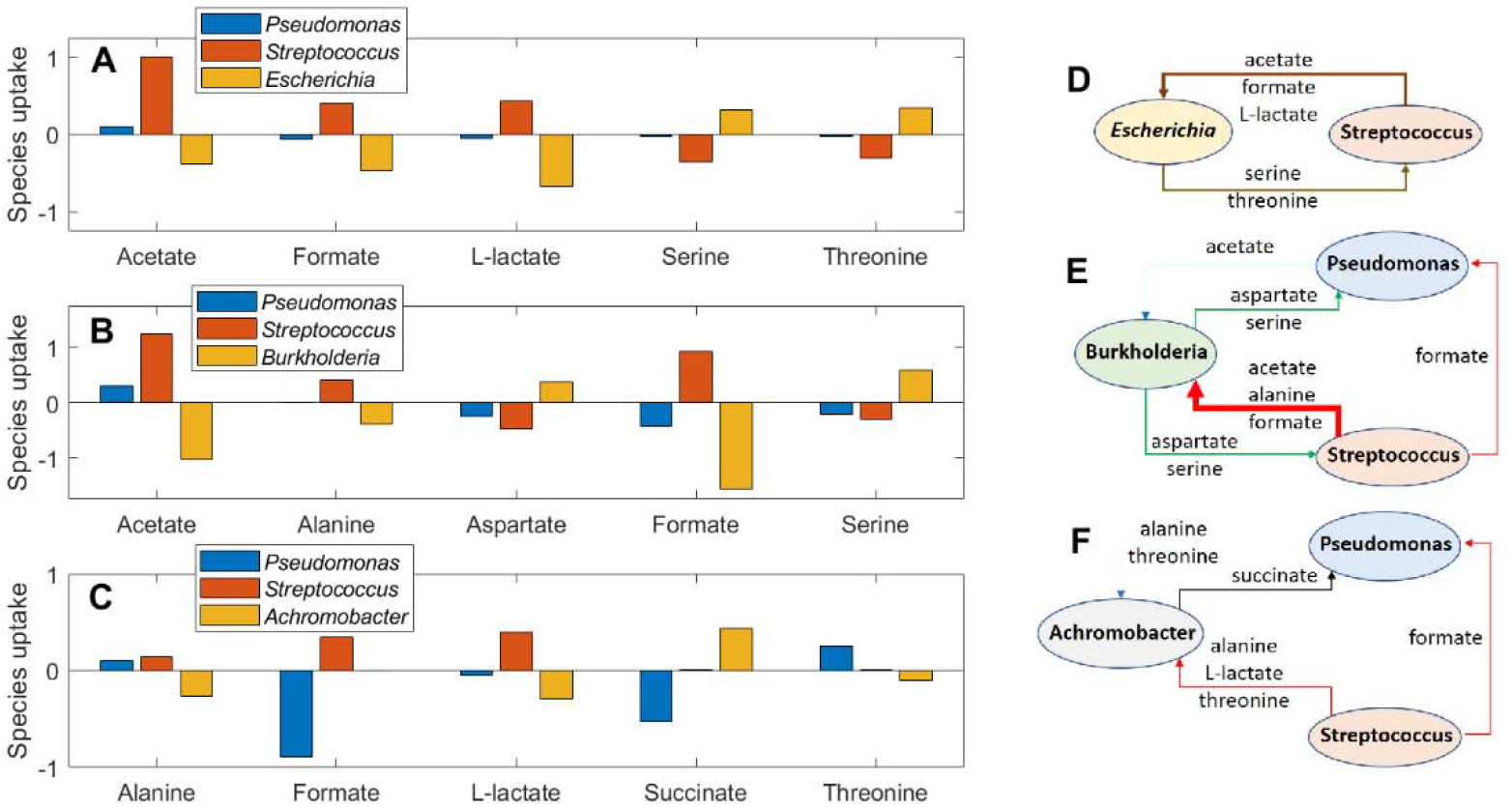
Predicted metabolite crossfeeding relationships for 15-species communities containing *Escherichia, Burkholderia* or *Achromobacter*. Negative rates denote metabolite uptake and positive rates denote metabolite secretion. The overall metabolite exchange rate from one species to another species was calculated by determining the minimum uptake or secretion rate for each exchanged metabolite and then summing these minimum rates over all exchanged metabolites. The arrow thickness is proportional to the overall metabolite exchange rate between the two species. (A) Average exchange rates of the five highest crossfed metabolites between the three most abundant species for 100 model ensemble simulations containing *Escherichia*. (B) Average exchange rates of the five highest crossfed metabolites between the three most abundant species for 100 model ensemble simulations containing *Burkholderia*. (C) Average exchange rates of the five highest crossfed metabolites between the three most abundant species for 100 model ensemble simulations containing *Achromobacter*. (D) Schematic representation of overall metabolite exchange rates for *Escherichia*-containing communities corresponding to Figure 7A. *Pseudomonas* was omitted due to its low exchange rates compared to the other two species. (E) Schematic representation of overall metabolite exchange rates for *Burkholderia-containing* communities corresponding to Figure 7B. (F) Schematic representation of overall metabolite exchange rates for *Achromobacter*-containing communities corresponding to Figure 7C.

The model ensemble also could reproduce variations in the second PC, which was heavily weighted by the three dominant genera and *Haemophilus*, for sufficiently large values of the first PC, which corresponded to relatively high *Pseudomonas* and low *Streptococcus* and *Prevotella*. By contrast, the model ensemble did not cover the patient samples in the lower left quadrant of the PC plot (Figure 6B). These samples were characterized by unusual combinations of relatively high *Prevotella, Haemophilus, Rothia* and/or *Fusobacterium* that the model could not reproduce in its present form. Of these 12 poorly modeled samples, *Prevotella* was highly represented in 8 samples. When the normalized reads of these 8 samples and their associated best-fit abundances were averaged, the models overpredicted *Pseudomonas, Streptococcus* and *Staphylococcus* at the expense of the less common genera (Figure 6C).

### The Community Model Predicts that Pathogen Dominance is Driven by Metabolite Cross-feeding

To investigate putative metabolic mechanisms by which pathogens may establish dominance in the CF lung, we used model predictions to quantify rates of metabolite cross-feeding between species. For each rare pathogen (*Escherichia, Burkholderia* and *Achromobacter*), 100 simulations performed with randomized community uptake rates were used to calculate average exchange rates of the five most significantly cross-fed metabolites between *Pseudomonas*, *Streptococcus* and the pathogen of interest. The overall metabolite exchange rate from one species to another species was calculated by determining the minimum uptake or secretion rate for each exchanged metabolite and then summing these minimum rates over all exchanged metabolites.

*Escherichia* was predicted to consume the organic acids acetate, formate and L-lactate produced by *Streptococcus*, while *Streptococcus* benefitted from the amino acids serine and threonine secreted by *Escherichia* (Figures 7A and 7D). Due to the existence of alternative optima with respect to the secretion products (49), L-lactate secretion was not predicted in *Streptococcus* monoculture even through the metabolic reconstruction supported L-lactate production (19) [www.vmh.life]. While *Streptococcus* strains are well known to product L-lactate as the primary product via homolactic fermentation (50, 51), we chose not to manually curate the metabolic reconstruction since *in silico* L-lactate synthesis was induced by the presence of other community members such as *Escherichia. Pseudomonas* was minimally involved in metabolite exchange due to its low average abundance (~1%) across the 100 simulations. Hence, our model suggested that organic acid cross-feeding could play a role in *Enterobacteriaceae* propagation in the CF lung.

More complex cross-feeding relationships were predicted for *Burkholderia*-containing communities that supported average *Pseudomonas* and *Streptococcus* abundances both exceeding 10%. The largest exchange rates were predicted for formate and acetate produced by *Streptococcus* and consumed by *Burkholderia* (Figures 7B and 7E). The two species also exchanged amino acids, with *Streptococcus* providing alanine to *Burkholderia*, and *Burkholderia* producing aspartate and serine for *Streptococcus*. *Burkholderia* provided the same two amino acids to *Pseudomonas* while receiving a small exchange of acetate in return. *Pseudomonas* also consumed formate secreted by *Streptococcus*. These model predictions suggested that acetate, formate and alanine produced by *Streptococcus* via heterolactic fermentation (50) could promote *Burkholderia* growth in vivo. Indeed, *in vitro* experiments have shown that mucin-degrading anaerobes such as *Streptococci* may promote the growth of CF pathogens such as *B. cenocepacia* by secreting acetate (43).

Compared to the other two pathogens, *Achromobacter* was predicted to be less efficient at cross-feeding having only small uptake rates of alanine, L-lactate and threonine secreted by the other two species. By contrast, *Pseudomonas* was predicted to benefit from relatively large uptake rates of formate produced by *Streptococcus* and succinate produced by *Achromobacter*. Collectively, these model predictions could help explain the enhanced ability of *Burkholderia* to dominate the simulated CF airway communities compared to *Achromobacter* (Figure 4) despite the single-species growth rates of the two species being similar (Figure 3).

Similar cross-feeding analyses were performed for 1000 simulations with randomized nutrient uptake rates in 14-species communities lacking *Escherichia*, *Burkholderia* and *Achromobacter*. To investigate the possibility of differential cross-feeding patterns, the simulations were split into 500 cases with the highest *Pseudomonas* abundances and 500 cases with the lowest *Pseudomonas* abundances (Figure 8A). For each set of 500 simulations, the average exchange rates of the five most significantly cross-fed metabolites between the four most abundant species (*Pseudomonas, Streptococcus, Prevotella* and *Staphylococcus*) were calculated. The overall metabolite exchange rate between any two species were calculated from the individual metabolite uptake and secretion rates as before.

**Figure 8.**
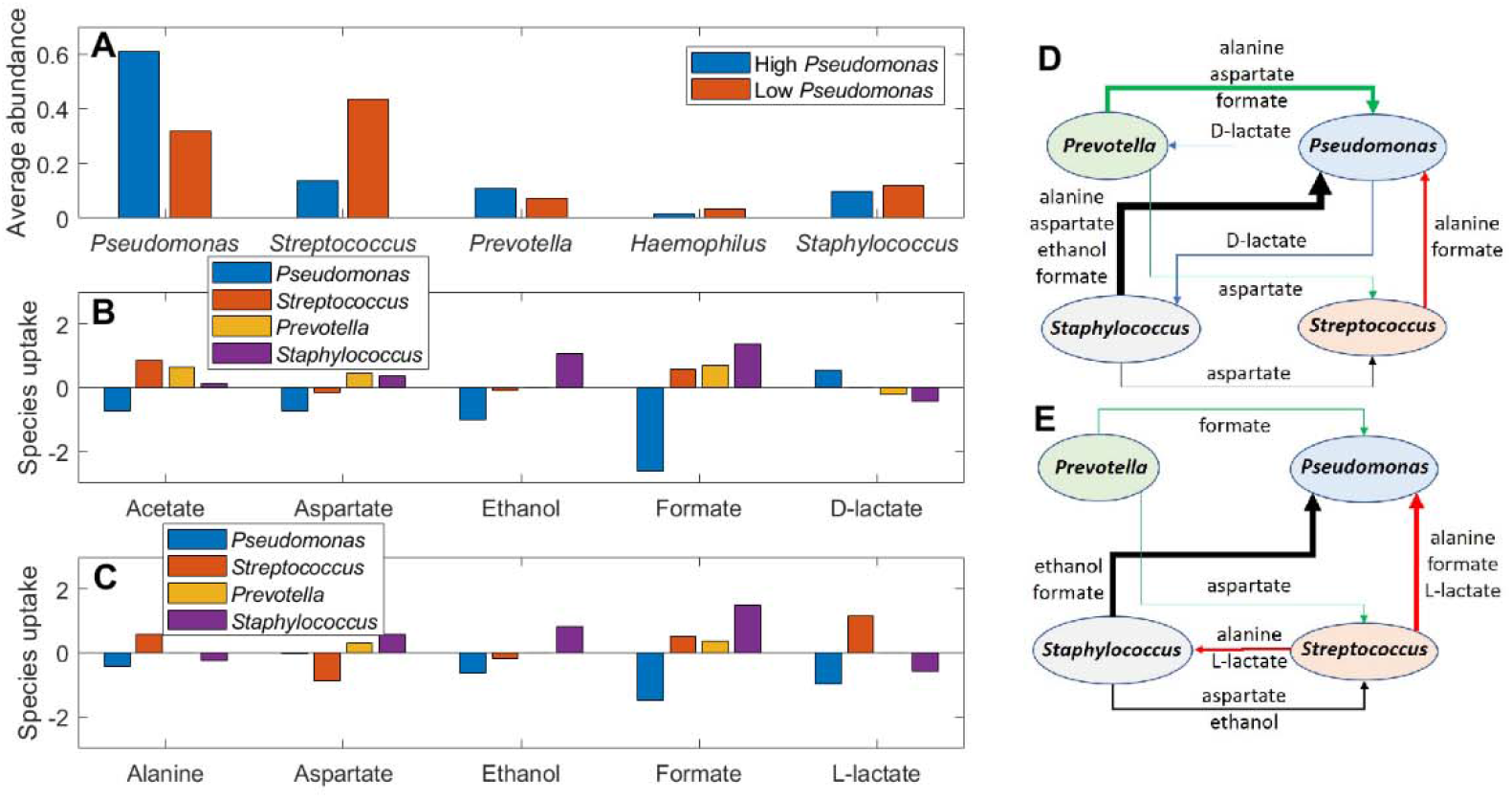
Predicted metabolite crossfeeding relationships for 14-species communities without *Escherichia*, *Burkholderia* and *Achromobacter*. 1000 model ensemble simulations were performed and split into 500 cases with relatively high *Pseudomonas* abundances and 500 cases with relatively low *Pseudomonas* abundances. (A) Average abundances of the five most highly represented species for the high and low *Pseudomonas* abundance cases. (B) Average exchange rates of the five highest crossfed metabolites between the four most abundant species for high *Pseudomonas* abundance cases. (C) Average exchange rates of the five highest crossfed metabolites between the four most abundant species the low *Pseudomonas* abundance cases. (D) Schematic representation of overall metabolite exchange rates for high *Pseudomonas* abundance cases corresponding to Figure 8B. (E) Schematic representation of overall metabolite exchange rates for low *Pseudomonas* abundance cases corresponding to Figure 8C.

When *Pseudomonas* abundances were predicted to be relatively high (average 61%), community interactions were dominated by *Pseudomonas* consumption of formate, ethanol, acetate and aspartate secreted by the other three species (Figure 8B). Formate cross-feeding was predicted to be particularly important, which was consistent with an *in vitro* study showing that expression of the *P. aeruginosa fdnH* gene encoding a formate dehydrogenase) was elevated in synthetic sputum medium compared to glucose minimal media (52). Similarly, the expression of the *P. aeruginosa adhA* (encoding an alcohol dehydrogenase) was elevated in patient-derived CF sputum compared to *in vitro* rich medium (53). Since *P. aeruginosa* strains have the capability to uptake both formate and ethanol (54, 55), these *in vitro* studies suggest that this cross-feeding mechanism could occur in CF airway communities. *Staphylococcus* was the major source of exchanged formate and ethanol (Figure 8D), a prediction consistent with studies showing that *P. aeruginosa* benefits from the presence of *S. aureus* (47, 56). Both alanine and aspartate have been shown to serve as preferred carbon sources for *P. aeruginosa* in a minimal medium supplemented with lyophilized CF sputum (52). However, the ensemble model did not predict exchange of L-lactate between *P. aeruginosa* and *S. aureus*, which differs from coculture experiments that mimic the CF lung environment (47). Strong interactions between *P. aeruginosa* and various *Streptococci* also have been reported (28), although the importance of metabolite cross-feeding in mediating these interactions remains incompletely understood (57). Finally, in the model *Pseudomonas* supplied small amounts of D-lactate for *Prevotella* and *Staphylococcus* consumption, a prediction consistent with an *in vitro* study showing *P. aeruginosa* anaerobic production of the LldA enzyme catalyzing D-lactate synthesis (58).

When *Pseudomonas* abundances were predicted to be relatively low (average 32%), metabolite cross-feeding remained dominated by *Pseudomonas* consumption of secreted byproducts and amino acids (Figure 8C). *Pseudomonas* was predicted to have high consumption rates of formate produced by all three other species and L-lactate synthesized only by *Streptococcus*, consistent with the ability of *S. salivarius* (59) and *P. aeruginosa* (47) to synthesize and consume L-lactate, respectively. Higher exchange rates between *Streptococcus* and *Staphylococcus* were predicted when *Pseudomonas* abundances were relatively low (Figure 8E). The two species cross-fed alanine and L-lactate produced by *Streptococcus*, and aspartate and ethanol secreted by *Staphylococcus*. Our predicted cross-feeding relationships in *Pseudomonas*- and *Streptococcus*-dominated communities could provide insights into CF disease progression, as high abundances of *Streptococcus* relative to *Pseudomonas* has been shown to correlate to higher diversity airway communities and improved CF clinical stability (28).

## Discussion

The airways of cystic fibrosis (CF) patients are commonly infected by complex communities of interacting bacteria, fungi and viruses which complicate disease assessment and treatment. The unique bacterial communities resident in individual patients can be longitudinally resolved to the genus level by applying 16S rRNA gene amplicon library sequencing to sputum and bronchoscopy samples (8). While 16S rRNA gene sequencing technology provides an unprecedented capability to identify bacterial pathogens in the CF lung, other analyses are required to understand how community members interact and how these interactions impede or promote disease progression. Metabolomics represents a powerful tool to interrogate the complex metabolic environment of the CF lung (60), but the number and depth of studies published to date has been limited. Metabolic modeling is a complementary tool for probing complex microbial communities and their interactions mediated through competition for host-derived nutrients and cross-feeding of secreted metabolites (11). Community metabolic models can provide information difficult to obtain by purely experimental means, such as the combined impact of nutrient environment and metabolic interactions on community composition. Metabolic models also can predict the rates of metabolite exchange between species and identify cross-feeding relationships difficult to delineate through metabolomic analyses.

We used 16S rRNA gene sequence data from three published studies (26-28) to construct and test a metabolic model for prediction of airway community compositions in adult CF patients. The assembled dataset consisted of 75 distinct samples from 46 patients who were judged to be stable or recovered from treatment in the original studies. Principal component analysis performed on 16S read data showed considerable heterogeneity of community composition across the 75 samples, including three patients infected with *Enterobacteriaceae, Burkholderia* and *Achromobacter* pathogens. Interestingly, each of these three patients was infected by only one of these “rare” pathogens, a characteristic we used to simplify our metabolic model simulations. The remaining 67 samples from 43 patients were largely dominated by *Pseudomonas* and/or *Streptococcus* but still exhibited substantial composition heterogeneity which provided a sufficiently-rich dataset to explore sample-to-sample variability.

The community metabolic model was constructed by ranking the identified taxa according to their total reads across the 75 samples and representing each taxonomic group with a single genome-scale metabolic reconstruction obtained from the AGORA database (www.vmh.life) (19). To limit model complexity, only the 17 top-ranked taxa (16 genera and 1 combined family/genus) were included. The resulting *in silico* community contained the most common CF pathogens (*Pseudomonas aeruginosa*, *Haemophilus influenzae, Staphylococcus aureus*), “rare” pathogens (*Escherichia coli, Burkholderia cepacian, Achromobacter xylosoxidans*), and 11 other species commonly observed in the CF sputum samples (e.g. *Prevotella melaninogenica, Rothia mucilaginosa, Fusobacterium nucleatum*). The 17 modeled taxa provided substantial coverage of the read data with an average coverage of 95.6+/-3.9% across the 75 samples. Because our *in silico* objective of growth rate maximization tends to produce low diversity communities dominated by ~5 species (39), the relatively low diversity of these adult CF lung samples made them particularly well suited for analysis through metabolic modeling as compared to considerably more diverse bacterial communities found elsewhere in the human body (*e.g*. intestinal tract (39, 61); chronic wounds (62)).

The community metabolic model required specification of host-derived nutrients that mimicked the CF lung environment in terms of the nutrients available, their allowed uptake rates across the community, and their allowed uptake rates by individual species. Given that the 17-species model contained 271 community uptake rates and a total of 2,378 species-specific uptake rates, a model tuning method was developed to manage the daunting complexity. A putative list of host-derived nutrients was compiled by starting with the synthetic sputum medium SCFM2 (63) and adding other nutrients either required for monoculture growth of at least one modeled species, measured in metabolomic analyses of CF sputum samples or identified through *in silico* analyses. The resulting 81 nutrients were separated into 14 distinct groups to facilitate tuning of nominal community uptake rates to qualitatively match average read data for the rare pathogen samples and the *Pseudomonas/Streptococcus-dominated* samples. This tuning process proved to be the bottleneck of model development even under the simplifying assumption that the species uptake rates were not limiting. A more streamlined and experimentally-driven tuning process would be facilitated by the availability of matched 16S and metabolomics data for large sets of CF sputum samples.

Despite the challenges associated with defining physiologically-relevant nutrient uptake rates, the community model was able to predict species abundance in qualitative agreement with average read data for *Enterobacteriaceae-, Burkholderia-, Achromobacter-* and *Pseudomonas/Streptococcus-dominated* samples. The modeling effort was simplified by omitting the other two rare pathogens when simulating the 3 *Enterobacteriaceae*-, 3 *Burkholderia*- and 2 *Achromobacter*-containing samples and omitting all three rare pathogens when simulating the other 67 samples, as justified through analysis of the 16S rRNA gene sequence data. The 15-species models used to simulate the rare pathogen-containing samples were able to reproduce dominance of the associated pathogen and, to a lesser extent, the abundances of less prevalent species. However, satisfactory prediction of the 2 *Achromobacter*-containing samples required the addition of four carbon sources (arabinose, fumarate, galactonate, xylose) which have not been measured in the CF lung to our knowledge. While there is some experimental evidence to support their inclusion, the need to add these four metabolites to elevate *in silico Achromobacter* growth could point to limitations of the modeled nutrients and their defined uptake rates.

The 14-species model used to simulate the rare pathogen-free samples predicted that *Pseudomonas* and *Streptococcus* would be the dominant genera, and that *Prevotella* and *Staphylococcus* also would be present in the community. These predictions provided qualitative agreement with the 16S rRNA gene sequence read data averaged across the 67 samples, although the predicted abundance of *Prevotella* was comparatively high and the predicted diversity was comparatively low. Given the uncertainty associated with identifying host-derived nutrients and translating these available nutrients into appropriate community uptake rates, we considered our predictions to provide satisfactory *in silico* recapitulation of measured community compositions across the set of four dominant CF pathogens.

A hallmark of CF lung infections is poorly understood differences in bacterial community compositions between patients and in longitudinal samples collected from a single patient (40). We performed simulations to test the hypothesis that these differences might be partially attributable to sample-to-sample variations in the nutrient environment in the CF lung. Nutrient variability was simulated by randomizing the community uptake rates around their nominal values found through manual model tuning. We performed 100 model ensemble simulations for each 15-species community containing a rare pathogen to determine if the associated patient samples could be well fit by a simulated sample. Using the least-squares difference between the measured reads and predicted abundances as the goodness-of-fit measure, we found that the model ensembles could satisfactorily reproduce the community compositions of the 8 rare pathogen-containing samples. The best-fit models tended to provide good predictions of rare pathogen reads due their relatively large values (average 65% across the 8 samples), while the accuracy of read predictions for less prevalent species was more variable.

Due to the availability of a much larger dataset of 67 patient samples, the rare pathogen-free model consisting of 14 species afforded an opportunity to investigate sample-to-sample heterogeneity in more depth. We performed 1000 model ensemble simulations with randomized nutrient uptake rates to find best-fit models. Patient samples with relatively high *Pseudomonas* reads tended to be well fit because the model predicted *Pseudomonas* dominance over a wide range of nutrient conditions. Less accurate but still satisfactory fits were obtained for patient samples with moderate *Pseudomonas* and relatively high *Streptococcus* reads. The model ensemble proved somewhat deficient in fitting samples with high reads of *Prevotella* or of the less common genera *Haemophilus, Rothia* and *Fusobacterium*. This deficiency could be attributable to the *in silico* lung environment not containing key nutrients and/or not specifying sufficiently large uptake rates of supplied nutrients to support high abundances of these genera.

The quality of sample fits also was correlated to the sample diversity, with the best fits having the lowest average diversity (inverse Simpson index of 0.10), moderate fits having an intermediate average diversity (inverse Simpson index of 0.18), and poor fits having the highest average diversity (inverse Simpson index of 0.23). For these three sets of samples, the best-fit models had average diversities of 0.10, 0.16 and 0.20, respectively. We believe that the lower predicted diversities were attributable to the modeling assumption that the CF lung community maximizes its collective growth rate. Using a community metabolic model of the human gut microbiota (39), we have shown that increased bacterial diversity (typically associated with health) can be achieved by simulating suboptimal growth rates under the hypothesis that disease progression correlates to a collective movement towards maximal growth. Therefore, the assumption of maximal community growth may inherently limit our ability to accurately reproduce more diverse samples, and rather simulate conditions associated with disease, such as dominance of a single pathogen.

By optimizing cross-feeding of secreted metabolites, the community model was able to predict the coexistence of multiple species at the maximal community growth rate rather than just predicting a monoculture of the single species with the highest monoculture growth rate. Because the SteadyCom method (25) used to formulate and solve the community model does not allow direct incorporation of mechanisms by which one species could inhibit the growth of another species other than by nutrient competition, the predicted community growth rate always was greater than the highest individual growth rate of the coexisting species. Consequently, the formulated model was incapable was capturing more complex interactions such as *Pseudomonas* secretion of diffusible toxins that inhibit the growth of other CF pathogens (64).

Despite this limitation, the community model could be analyzed to understand the putative role of metabolite cross-feeding in shaping community composition. The model predicted that the rare pathogens *Escherichia* and *Burkholderia* were particularly efficient cross-feeders, using acetate, formate and other secreted metabolites to establish dominance over less harmful bacteria. By contrast, the model predicted *Achromobacter* to be substantially less adept at exploiting secreted metabolites for growth enhancement. While we were able to simulate *Achromobacter* dominance through addition of four carbon sources possibly present in the CF lung, the model suggested that other non-modeled mechanisms may be involved in promoting *Achromobacter* expansion. One possibility is that *Achromobacter* utilizes its ability to form multispecies biofilms (44, 65) to establish favorable metabolic niches for enhanced growth.

In the absence of the three rare pathogens, the model predicted that *Pseudomonas* would be the primary beneficiary of cross-fed metabolites including acetate, alanine and L-lactate from *Streptococcus* and aspartate, ethanol and formate from *Staphylococcus*. These complex cross-feeding relationships were an emergent property of the community model that could not predicted from monoculture simulations and are consistent with published experimental data presented above. For example, the single-species models predicted that acetate, CO_2_ and formate would be the primary secreted byproducts yet the community model also cross-fed ethanol, D-lactate, L-lactate and succinate which were not predicted to be secreted in any monoculture simulation. We hypothesized that model ensemble simulations with relatively high and low *Pseudomonas* abundances would show differential cross-feeding patterns. While some of the specific cross-fed metabolites changed between the two cases, cross-feeding from *Streptococcus* and *Staphylococcus* to *Pseudomonas* remained the dominant feature of the simulated communities. In our assimilated dataset of 75 patient samples, *Pseudomonas* reads were above 10% in 55 samples and above 50% in 35 samples. Our model predictions provide putative metabolic mechanisms that may help explain why *Pseudomonas* so efficiently colonizes the adult CF lung and why *Pseudomonas* commonly establishes dominance over other species once colonized.

Our community metabolic model generated several predictions that could be tested experimentally with an appropriately designed *in vitro* community. For example, a 5-species *in vitro* system consisting of *Pseudomonas aeruginosa, Streptococcus sanguinis, Prevotella melaninogenica, Haemophilus influenzae* and *Staphylococcus aureus* would provide substantial coverage of our 16S rRNA gene sequencing data as the five genera account for 81% of reads across the 75 samples and greater than 75% of reads in 56 samples. Specific model predictions that could be tested *in vitro* include the variability of community compositions by changing nutrient levels in a synthetic CF medium, and the cross-feeding of specific metabolites by genetically altering the secretion and/or uptake capabilities of these metabolites in the relevant species. The availability of such *in vitro* data linking the nutrient environment, cross-feeding mechanisms and community composition would allow direct testing of a simplified 5-species model and facilitate the development of improved community models for the analysis of CF sputum samples.

## Materials and Methods

### Patient Data

CF airway community composition data was obtained from three published studies in which patient sputum samples were subjected to 16S rRNA gene amplicon library sequencing (26-28). The assimilated dataset contained 75 distinct samples from 46 patients who were clinically stable or recovered from treatment for an exacerbation event. Additional samples from these three studies corresponding to exacerbation or antibiotic treatment were not included in the modeled dataset to avoid the complications of predicting these events. The top 72 taxonomic groups (typically genera) accounted for over 99.8% of total reads across the 75 samples (Figure 1A; Table S1). To limit complexity, the community metabolic model described below was limited to 17 taxonomic groups that accounted for 95.6% of total reads (Figure 1B; Table S2). Reads from the family *Enterobacteriaceae* and the genus *Escherichia* were combined and represented as a single genus. To allow direct comparison with the species abundances predicted by the model, the reads for each sample were normalized over the 17 modeled genera to sum to unity (Table S3).

### Community Metabolic Model

For simplicity, each genus was represented by a single species commonly observed in CF airway communities (1, 6-9, 66), although we note that genera such as *Streptococcus* (28) can have considerably diversity with respect to species representation. As mentioned above, the combined *Enterobacteriaceae/Escherichia* taxonomic group was represented by the single species *Escherichia coli*. A genome-scale metabolic reconstruction for each species (Figure 1C) was obtained from a large database of AGORA models (19) (www.vmh.life). Table 1 lists the representative strain used for each genus, the normalized reads fractionally associated with each genus averaged across the 75 samples (also shown in Figure 1B), and the number of samples for which the normalized reads exceeded 1%. The community model accounted for 13,845 genes, 19,034 metabolites and 22,412 reactions within the 17 species as well as 271 uptake and secretion reactions for the extracellular space shared by the species.

The genera *Pseudomonas, Streptococcus* and *Prevotella* dominated most communities, both in terms of average reads for individual samples and the number of samples in which they exceeded 1%. Interestingly, *Enterobacteriaceae/Escherichia, Burkholderia* and *Achromobacter* exceeded 0.1% in only single patients represented by 3, 3, and 2 samples, respectively. Moreover, no patients were infected by more than one of these “rare” pathogens, as the maximum reads of the other two pathogens never exceeded 0.1% in these 8 samples. Therefore, for modeling purposes the 75 samples were partitioned into: 3 *Enterobacteriaceae/Escherichia-containing* samples with *Burkholderia* and *Achromobacter* absent; 3 Burkholderia-containing samples with *Enterobacteriaceae/Escherichia* and *Achromobacter* absent; 2 Achromobacter-containing samples with *Enterobacteriaceae/Escherichia* and *Burkholderia* absent; and 67 samples with all three rare pathogens absent.

### Model Tuning and Simulation

The nutrient environment in the CF lung is complex and expected to vary between patients as well as between longitudinal samples for individual patients depending on disease state. While metabolomic analyses have been performed on CF sputum and bronchoscopy samples (40, 60, 66, 67), these studies were insufficient to define supplied nutrients for the metabolic model due to their limited metabolite coverage. Furthermore, we found that based on our model, the synthetic sputum medium SCFM2 used in previous *in vitro* CF microbiota studies (63, 68) would not support growth of any of the 17 modeled species due to the lack of ions (Co^2+^, Cu^2+^, Mn^2+^, Zn^2+^), amino acids (asparagine, glutamine) and other metabolites (see below) essential for growth. While the medium likely would contain trace amounts of the missing ions, the requirement of these other metabolites for growth suggests limitations for the AGORA metabolic models with respect to biosynthetic pathways leading to biomass formation. Given the semi-curated nature of the AGORA models (19), such discrepancies were expected and had to be addressed by adding the missing essential metabolites to the modeled medium. A final complication was that the community model required specification of nutrient uptake rates, which were unknown even if medium component concentrations were specified due to the lack of species-dependent uptake kinetics for each nutrient. Because such uptake information is rarely available even for highly studied model organisms such as *Escherichia coli* (69), a simplified approach was used to define nutrient uptake rates for the community model.

Supplied nutrients in the community model were defined by starting with the SCFM2 medium and adding the four ions and two amino acids listed above. We found that each species required additional metabolites in the medium to support biomass formation. These 29 additional metabolites were identified and added to the modeled medium such that all 17 species were capable of monoculture growth (see Table S4). For example, the *P. aeruginosa* model required addition of uracil and menaquinone 7, while *in vitro* experiments have shown that these metabolites are synthesized *de novo* and not required in the medium (63). Next, we added four carbon sources (fructose, maltose, maltotriose, pyruvate) and 8 other metabolites (adenosine, cytidine, glycerol, guanosine, hexadecanoate, inosine, octadecenoate, uridine) measured in the CF lung (67) and the terminal electron acceptor O_2_ to simulate aerobic respiration. Finally, we added four additional carbon sources (arabinose, fumarate, galactonate, xylose) that increased *in silico Achromobacter* growth such that *Achromobacter* would be competitive with other species when it was present in the community. While these carbon sources were identified *in silico*, there is experimental evidence to support their inclusion in the simulated CF lung environment. Fumarate has been shown to be elevated in sputum samples from young CF patients (70). Arabinose and xylose are constituents of extracellular polymer substance (EPS) produced by common human pathogens including the modeled genera *Pseudomonas, Staphylococcus* and *Escherichia* (71), suggesting their possible presence in the CF lung. Pathogenic *Achromobacter* strains isolated from CF patients has been shown to grow on galactonate as a sole carbon source (72), supporting the hypothesis that *Achromobacter* has evolved to utilize galactonate available in the CF lung.

The community uptake rates of the 86 supplied nutrients were tuned by trial-and-error to produce species abundances in approximate agreement with the average reads listed in Table 1, which were derived from actual patient samples. To reduce the number of adjustable rates, the nutrients were grouped together and a single uptake rate was used for each group. These 14 groups were defined as: (1) 16 common metals and ions; (2) 29 essential growth metabolites; (3) 8 CF lung metabolites; (4) 19 amino acids; (5) the amino acids alanine and valine, which have been reported to be elevated in the CF lung compared to other amino acids (67); (6)-(11) each of the 6 carbon sources available in the CF lung; (12) O_2_; (13) NO_3_; and (14) 4 *Achromobacter-related* carbon sources. The 86 nutrients and their nominal community uptake rates determined through this tuning procedure are listed in Table S4 and depicted graphically in Figure 1D.

Because these nutrient uptakes rates were derived for the entire patient population and not an individual patient sample, a different strategy was used to simulate sample-to-sample heterogeneity based on the hypothesis that differences in nutrient availability could account for heterogeneity in measured reads. Individual patient samples were simulated by randomly perturbing the community uptake rate for each of the 14 nutrient groups listed above between 33% and 300% of its nominal value. Uniformly distributed random numbers were generated for each group such that the number of cases with the uptake rates in the range [33%-100%) and [100%-300%] were statistically equal. The bounds used for the uptake rate of each metabolite also are listed in Table S4.

### Community Simulations

We used the SteadyCom method (25) to perform community simulations as detailed in our previous study on the human gut microbiota (39). SteadyCom performs community flux balance analysis by computing the relative abundance of each species for maximal community growth while ensuring that all metabolites are properly balanced within each species and across the community. Each species model used a non-growth associated ATP maintenance (ATPM) value of 5 mmol/gDW/h, which is within the range reported for curated bacterial reconstructions. Cross-feeding of all 21 amino acids and 8 common metabolic byproducts (acetate, CO_2_, ethanol, formate, H_2_, D-lactate, L-lactate, succinate) was promoted by increasing the maximum nutrient uptake rates of these nutrients in each species model to 2.5 and 5 mmol/gDW/h, respectively. Outputs of each SteadyCom simulation included the community growth rate, the abundance of each species, and species-dependent uptake and secretion rates of each extracellular metabolite. The nominal nutrient uptake rates produced a single community not directly comparable to any single patient sample (Figure 1E), while each set of randomized uptake rates produced a unique community that was interpreted as a prediction of an individual patient sample (Figure 1F).

## Data availability

All data used for metabolic model development and testing is provided in the Supplemental Material.

## Acknowledgements

The authors wish to acknowledge the NIH grants R37 AI83256-06 (GAO) and T32-AI007519 (GO) for partial support of this research.

## Supplementary Materials

Table S1. 16S sequencing reads for the top 72 taxonomic groups assembled from three published CF studies.

Table S2. 16S sequencing reads for the top 17 taxonomic groups assembled from three published CF studies.

Table S3. Normalized 16S sequencing reads for the top 17 taxonomic groups assembled from three published CF studies.

Table S4. Minimum, nominal and maximum community uptake rates for supplied nutrients.

Table S5. Principal component analysis of normalized read dataset containing all 75 samples.

Table S6. Principal component analysis of normalized read data excluding 8 samples containing *Enterobacteriaceae/Escherichia, Burkholderia* and *Achromobacter*.

Table S7. Comparison of normalized reads and model predicted abundances for 8 patients samples containing the pathogen *Enterobacteriaceae/Escherichia, Burkholderia* and *Achromobacter*.

Table S8. Comparison of normalized reads and model predicted abundances for 9 representative patient samples not containing the pathogens *Enterobacteriaceae/Escherichia, Burkholderia* or *Achromobacter*.

